# A method for correcting background non-uniformity while preserving specimen-level information in Fourier ptychographic microscopy images

**DOI:** 10.64898/2026.10.21.760683

**Authors:** L. Copeland, K. Polland, L.D. Walker, W.B. Amos, K.J. Baxter, M. Shaw, G. McConnell

**Author notes:** Laura Copeland, Kay Polland, and Lewis Walker are joint first authors.

## Abstract

Fourier ptychographic microscopy (FPM) enables high resolution imaging across large fields of view, but reconstructed images often exhibit background non-uniformity that interferes with quantitative image analysis. Here, a post-reconstruction background correction approach is presented that identifies specimen-containing regions and corrects only the surrounding background, leaving specimen pixel intensities unchanged. The method was effective across amplitude and phase reconstructions of multiple biological specimen types with no loss in image resolution. The method also improved object detection compared with analysis of the uncorrected image data. FPM background correction reduced residual background non-uniformity to 0.61% with only 0.04% structural error, compared with 6.13% structural error for pseudo flat field correction. These results demonstrate that FPM background correction can support more accurate quantitative analysis of FPM image data while preserving specimen information and image resolution, enabling more reliable segmentation and measurement across large fields of view.

## 1. Introduction

Fourier ptychographic microscopy (FPM) is a computational imaging method that combines a sequence of low resolution images acquired under spatially varying illumination angles to reconstruct a high resolution image over a large field of view.^1,2^ By synthesising a numerical aperture greater than that available from the objective lens alone, FPM can overcome the conventional trade-off between resolution and imaging area, known as the space-bandwidth product.^3^ The resulting amplitude and phase reconstructions have made FPM attractive for applications requiring quantitative or high-throughput imaging across extended specimens, including cell and tissue imaging, pathology, and other forms of biological microscopy.^2,4–8^ An important consequence of this combination of large imaging area and computational reconstruction, however, is that non-uniformity in illumination and reconstruction can become apparent across the reconstructed field.

Background non-uniformity and artefacts can arise at several stages of the FPM imaging and reconstruction process. FPM is sensitive to residual background signals and noise in the acquired image sequence, with inadequate removal of dark-field background demonstrated to produce substantial artefacts in reconstructed images.^9,10^ Variation in illumination intensity between light emitting diodes (LEDs), together with imperfections in the optical system and reconstruction process, can introduce further variation into the reconstructed image.^9–13^ The reconstructed image in FPM is produced patch-wise due to limited spatial coherence of the illumination source, and because of the large size of the raw datasets. In practice, these effects can produce intensity gradients, step changes in background intensity, band-like structure, and localised regions of brighter or darker background appearing similar to stitching and tiling artefacts.^9,10,12^ However, these are not necessarily present as identical artefacts in the amplitude and phase image outputs. Such artefacts can extend across substantial regions of an FPM reconstruction and may co-exist within the same image, obscuring weak specimen features, reducing local contrast, and making quantitative analysis difficult.

Background correction is well established in microscopy^14–17^. Conventional flat-field correction relies on measurement of an appropriate reference field^15,16^, while approaches such as BaSiC estimate background non-uniformity from collections of images.^17^ In FPM, background non-uniformity has previously been addressed through adaptive separation of background interference^10^ and suppression of background signal and noise in dark-field acquisitions.^9^ However, these approaches are applied to either the raw image sequence via preprocessing or during reconstruction, rather than on the reconstructed output, and they do not remove artefacts associated with patch-wise reconstruction. There is therefore a need for a background correction method that operates directly on FPM images while preserving specimen-associated pixel values.

Here, a post-reconstruction background correction method for FPM images is presented in which specimen-containing regions are identified from local image structure and protected from subsequent modification, allowing background non-uniformities to be removed from the surrounding image background. The approach is evaluated using both amplitude and phase contrast reconstructions, for multiple biological specimen types, and for different patterns of background non-uniformity. The consequences for downstream quantitative analysis are examined using automated object detection, while image decorrelation analysis is used to determine whether improved background uniformity is achieved without loss of image resolution. The method is also compared quantitatively with pseudo flat field correction, examining both residual background non-uniformity and changes to specimen information. Together, these experiments assess whether background non-uniformity in reconstructed FPM images can be reduced sufficiently to improve quantitative image analysis without altering the specimen-level information.

## 2. Methods

### 2.1 FPM background correction

The post-reconstruction background correction pipeline was implemented in Python. The method was designed to reduce background non-uniformity while preserving specimen-associated pixel values. Rather than applying a correction across the complete image, the approach first identifies regions containing specimen features and protects these regions from subsequent modification. Background correction was then restricted to pixels outside the protected mask. The overall workflow, comprising multiscale Gaussian filtering, specimen detection and mask generation, and selective background replacement, is illustrated in Figure 1.

**Figure 1.**
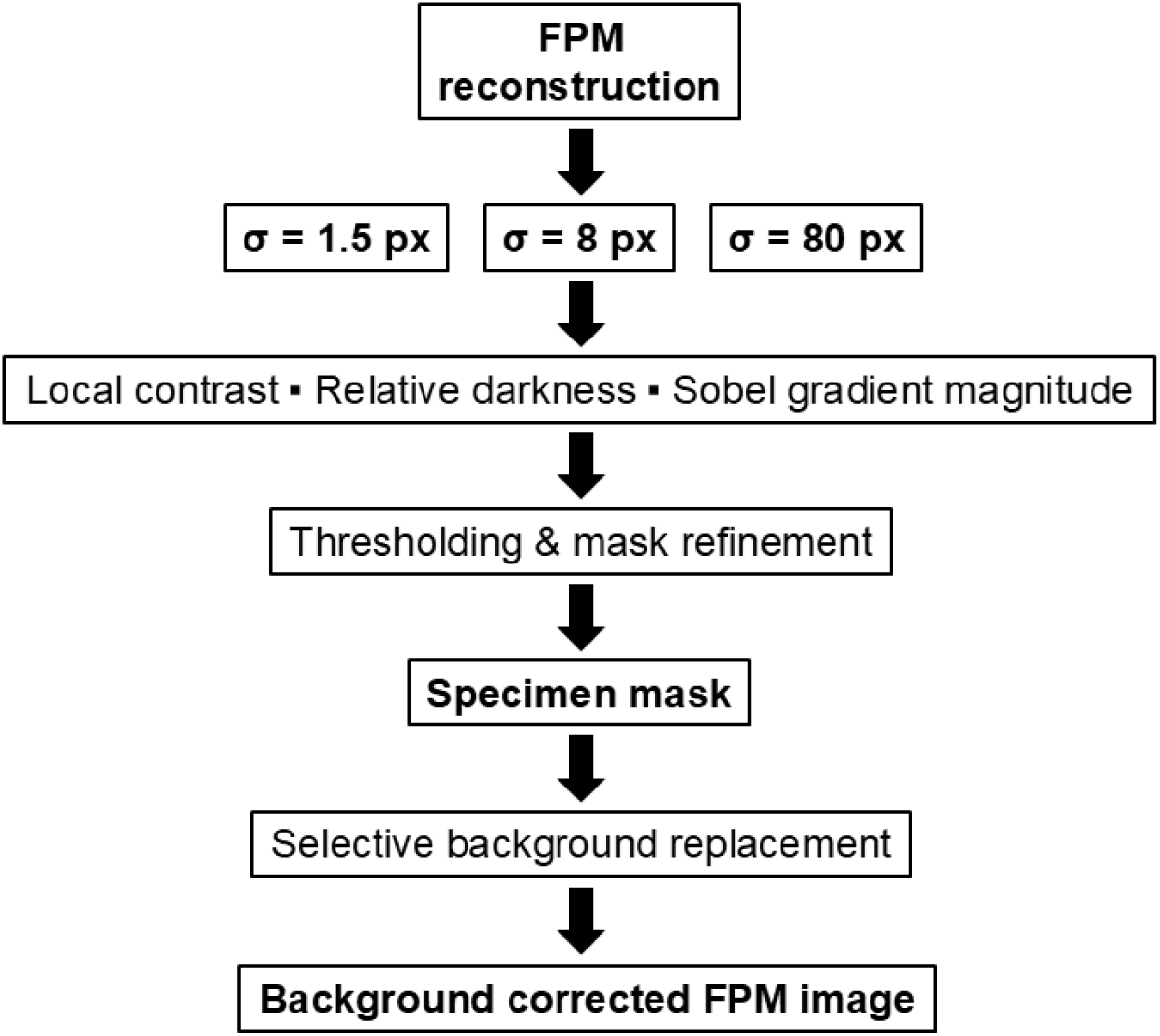
Workflow of the FPM background correction method. An FPM amplitude or phase reconstruction is Gaussian-filtered at three spatial scales (σ = 1.5 pixels, 8 pixels, and 80 pixels). Local contrast, relative darkness, and Sobel gradient magnitude are calculated from the filtered images to identify regions of the image containing the specimen. Thresholding and mask refinement are then applied to generate a binary specimen mask. Pixels outside the mask are replaced with the median background intensity, while protected specimen pixels retain their original values, producing a background corrected FPM image.

The software implementation used open-source Python libraries including NumPy^18^, SciPy^19^, imageio, Pillow, and tifffile for numerical operations, image filtering, morphological processing, and file input/output. The numerical datatype of the input image was preserved when saved in a compatible image format.

Execution time was measured using Python’s time.perf_counter() around the complete processing workflow, including image loading, specimen detection, background correction, output and diagnostic file saving, and verification of the saved image.

In this work, an HP EliteBook 830 computer with 1.6 GHz Intel® Core™ Ultra 5 processor, with 16 GB RAM and Microsoft Windows 11 Enterprise was used.

#### 2.1.1 Image representation and preprocessing

Reconstructed amplitude and phase images were converted internally to 32-bit floating-point representation for processing without rescaling their intensity values. As all FPM reconstructions analysed in this study were greyscale, the input image was used directly for specimen detection. The detection procedure was applied to the supplied amplitude or phase reconstruction, with the resulting mask used to identify pixels protected from subsequent background correction.

To distinguish the specimen from variations in the reconstructed background, the reconstructed amplitude or phase image was filtered at three different spatial scales using Gaussian filters with σ=1.5 pixels, 8 pixels, and 80 pixels. This generated high, intermediate, and low resolution representations, respectively. These filtered representations were used exclusively for specimen detection and were not used to reconstruct or modify specimen pixel values. The Gaussian filter widths therefore determined the spatial scales contributing to mask generation rather than the spatial resolution of the corrected image. All protected pixels were retained from the original, unfiltered reconstruction.

Local image detail was estimated from the absolute difference between the high resolution and intermediate resolution images, while relative darkness was determined from the difference between the low resolution and high resolution images. This multi-scale representation suppresses slowly varying background intensity changes during specimen detection while retaining local features associated with the specimen.

#### 2.1.2 Specimen detection and mask generation

Regions of the image containing the specimen were identified using a combination of local contrast, relative darkness, and image-gradient information. Local contrast was obtained from the difference between the high resolution and intermediate resolution images, while relative darkness identified structures that were locally darker than the low resolution image background. Image edges were detected by applying Sobel filters in the horizontal and vertical directions to the high resolution image and calculating the corresponding gradient magnitude.

To provide thresholds that were insensitive to the absolute intensity range of the input image, each of these feature images was converted to a robust z-score using the median and median absolute deviation. Pixels exceeding robust z-score thresholds of 3.5 for local contrast, 3 for relative darkness, or 4 for gradient magnitude were classified as candidate specimen pixels. The three resulting masks were combined using a logical OR operation so that a pixel identified by any of the detection criteria was included in the initial specimen mask.

Morphological processing was subsequently used to generate a continuous protected specimen mask. Binary closing was performed for two iterations using a 3 x 3 cross-shaped structuring element, followed by filling of internal holes. Connected foreground components containing fewer than 30 pixels were then removed. The resulting mask was dilated by four iterations using the same structuring element to provide an additional protected region around detected specimen boundaries. This dilation reduced the likelihood that specimen edges or immediately adjacent intensity features were inadvertently modified during background correction.

The final binary mask separated the image into protected specimen-containing regions and unprotected background regions. Only pixels classified as background were modified during the subsequent correction step.

#### 2.1.3 Background correction

Background correction was performed using only pixels outside the protected specimen mask. The target background intensity was calculated as the median intensity of all unprotected pixels. The median provided a robust estimate of the central background intensity while reducing sensitivity to residual high or low intensity features that were not included in the specimen mask.

For the correction used in this study, all unprotected pixels were replaced with this median background value. Protected specimen pixels were not corrected, filtered, or interpolated: these were instead copied directly from the corresponding positions in the original image. Consequently, the correction removed variation from regions identified as background without changing the protected signal from the specimen. No attempt was made to estimate or subtract background contributions within the protected specimen regions. This was a deliberate design choice, as correction of specimen intensities would require assumptions about the underlying background signal and could introduce changes to the information being measured. The method therefore prioritised preservation of the original specimen pixel values rather than correction of background contributions potentially present within the specimen itself.

Pixels within the protected mask were explicitly copied from the original working image into the corrected image. An element-wise comparison of the protected pixels in the original and corrected arrays was then made. Processing was terminated if any difference was detected, providing an internal check that the correction had not modified protected specimen pixels. The corrected image was subsequently converted back to the data type of the input image.

### 2.2. Specimen preparation and imaging

Several different specimens were used to evaluate the method. All specimens were mounted between a Type 1.5 coverslip (48393-172, VWR, US) and a microscope slide (TT-7041I-8217-001, Epredia, US).

*Arabidopsis thaliana* Col-0 (N1092, Nottingham Arabidopsis Stock Centre, UK) seeds were used. The seeds were sterilised in a bleach-based solution described in Supplementary Material and stored in the fridge for 2 days. The seeds were then plated onto ½ Murashige and Skoog (MS) agar plates^20^ and grown at 22°C in a 12-hour light-dark cycle. The roots were separated from the rest of the plant and mounted in 50 μL deionised water between slide and coverslip.

Human palatine tonsil samples were collected from the Royal Hospital for Children, Glasgow, UK. The ethics were approved by Biorepository Management Committee of NHS Greater Glasgow and Clyde, UK (NHS Biorep 548). Specimens were then transported to the University of Strathclyde in a sterile saline solution (0.9% Sodium chloride, Baxter Healthcare Ltd, UK) in less than 2 hours post excision. Whole fresh tonsils were frozen in optical cutting temperature (OCT) embedding matrix for frozen tissue (KMA-0100-00A, Cell Path, UK). Whole tonsils were sectioned to a thickness of 10 µm using a microtome (CM1950, Leica Microsystems, Germany) and specimens were mounted on microscope slides (Z692255, Sigma-Aldrich, USA). This thickness was chosen to reduce tissue folding and tearing. Tissue sections were fixed in 4% paraformaldehyde followed by a series of 3 washes for 5 min each using 1X phosphate buffered saline (PBS) (10010023, ThermoFisher Scientific, USA). Tonsils were covered with Haematoxylin mayer (HX29525075, Sigma-Aldrich, USA) for 2.5 minutes then rinsed twice with dH_2_O. Bluing reagent (AB245880, Abcam, UK) was then added to each section for 10-15 seconds then rinsed twice with dH_2_O. Absolute alcohol was added then immediately tipped off slides. Eosin y (318906, Sigma-Aldrich, USA) solution was added to each section for 1.5 minutes then rinsed with absolute alcohol 4 times. The specimens were air-dried and mounted under Type 1.5 coverslips using clear nail polish to set down the edges of the coverslips.

*Nicotiana benthamiana* seeds were sterilised using a bleach-based solution as described in the Supplementary Material and stored in the fridge for 2 days. The seeds were then plated onto ½ MS plates^20^ and grown at 22°C in a 12-hour light-dark cycle. The roots were separated from the rest of the plant and mounted in 50 μL deionised water between a slide and coverslip.

*Spirogyra varians* (LZA075, Blades Biological Ltd, UK) was grown using 3N-BBM+V (Culture Collection of Algae & Protozoa, UK). The algae were mounted using 50 μL deionised water between slide and coverslip.

*Tetrahymena pyriformis* was prepared for imaging using a wet silver impregnation method^21^ as described in detail by Mackinnon and Hawes^22^. Briefly, ciliates were chemically fixed, embedded in saline gelatin, and impregnated with 3% aqueous silver nitrate. Silver reduction was induced using a germicidal ultraviolet lamp until the gelatin surrounding the specimens developed a brown colour, after which the preparation was mounted between slide and coverslip.

Silicon dioxide microspheres of 5 µm diameter (44054, Merck, UK) were diluted 1:100 (v/v) in dH_2_O. To prepare the specimen for imaging, 10 µl of the suspension was pipetted on microscope slides (Z692255, Sigma-Aldrich, USA) and left to air dry. The specimen was mounted in Histomount (NAT1310, Scientific Laboratory supplies, UK) and a coverslip added before setting.

Specimens were imaged using an existing FPM system, with details of the microscope published previously^7^. Briefly, the system included a horizontal optomechanical imaging configuration with a vertically mounted planar 16×16 LED array with a 10 mm pitch (WS2812B, BTF lighting). The LED array was positioned 121 mm from the specimen, which was mounted on a two-axis translation stage (XYF1, Thorlabs) for lateral positioning and a single-axis translation stage (PT1, Thorlabs) for focusing along the optical axis. Brightfield and dark-field images were acquired under sequential multi-angle illumination at a wavelength of 636 nm from 177 LEDs arranged in a circular pattern. Each dataset consisted of 9 brightfield and 168 dark-field images acquired using a monochrome CMOS camera (UI-3060CP-M-GL R.2, iDS). Images were formed on the camera sensor using an infinity-corrected imaging system including a 4x/0.13 NA Plan Fluor objective lens with a 17.2 mm working distance (Nikon) and a 200 mm tube lens (TTL200-A, Thorlabs). LED illumination and camera hardware triggering were controlled and synchronised using open-source software running on a Raspberry Pi 4 model B^23^. Raw 8-bit images were acquired and stored on an external PC using ThorCam, with individual FPM image tiles subsequently reconstructed in MATLAB using the sequential quasi-Newton phase retrieval algorithm on open-source software.^24^ Lastly, reconstructed tiles were stitched together using FIJI to produce full field of view high resolution amplitude and phase images.^25^

### 2.3 Assessment of background correction

The performance of the method was assessed by comparing reconstructed FPM images before and after background correction. Raw and corrected images were displayed using identical intensity limits to permit direct visual comparison of background non-uniformity. Pseudocolour representations using the ‘16 Colors’ look-up table (LUT) in FIJI were also generated with the same display limits before and after correction to enhance the visibility of relatively small variations in background intensity.

The spatial distribution of intensity changes introduced by the correction was assessed using absolute pixel-wise difference images calculated between the raw and corrected reconstructions. These images were used to confirm that intensity changes were predominantly confined to regions identified as background, with specimen-containing regions remaining unchanged.

To determine that background correction did not degrade image resolution, resolution was measured using the Image Decorrelation Analysis plugin^26^ for FIJI with default settings. Three 2000 pixel x 2000 pixel regions of interest (ROIs) were selected from images of tonsil tissue, *S. varians*, *N. benthamiana*, and *T. pyriformis*. ROIs were chosen from different regions of the field of view with each ROI selected to include specimen information rather than background alone. The same ROIs were analysed in the corresponding raw and FPM background corrected reconstructions. Differences in decorrelation-based resolution before and after background correction were assessed using a two-tailed paired *t*-test, with *P* < 0.05 considered statistically significant.

Spatial frequency content was also assessed qualitatively using 2D fast Fourier transforms (FFTs) of the original and background corrected *N. benthamiana* amplitude and *T. pyriformis* phase reconstructions. Corresponding Fourier spectra were compared visually to assess whether background correction altered the overall distribution and extent of spatial frequency content.

### 2.4 Assessment of object detection before and after FPM background correction

The effect of FPM background correction on automated object detection was assessed using amplitude and phase reconstructions of 5 µm silica beads mounted in Histomount (NAT1310, Scientific Laboratory supplies, UK). Raw and background-corrected reconstructions were independently thresholded in FIJI using the Otsu method.^27^ The resulting binary images were analysed using the Analyze Particles tool with a size range of 0.8 µm²-Infinity and a circularity range of 0-1, with no additional options selected. Identical ROIs, each containing three visually identifiable silica beads, were analysed in the corresponding raw and background-corrected images. The number of objects detected within each ROI before and after background correction was recorded.

### 2.5 Comparison of the FPM background correction with pseudo flat field correction

The FPM background correction method was compared with pseudo flat field correction from the BioVoxxel Toolbox (v2.6.0a).^28^

A raw FPM amplitude image was corrected using the FPM background correction method and using pseudo flat field correction with blur radii of r=50, r=150, r=250 and r=350 pixels. Specimen and background regions were identified automatically from the raw image, and the same masks were used for all corrected images. The raw image was smoothed with a Gaussian filter (σ = 4 pixels) and the specimen was identified using Otsu thresholding. Small objects and holes were removed and morphological closing was applied. The background mask excluded the specimen together with a halo of 30 pixels.

Background non-uniformity was measured using non-overlapping 64 pixel x 64 pixel tiles. Tiles containing at least 90% background were included and the median background intensity was calculated for each tile. Residual background non-uniformity was calculated as the coefficient of variation of the tile medians.

To assess changes to the specimen, pixel intensities within the specimen mask were compared with the corresponding pixels in the raw image. A linear fit was used to allow for changes in overall intensity and offset, and the root-mean-square error of the residuals was calculated. This was normalised to the 5^th^ to 95th percentile intensity range of the raw specimen and expressed as a percentage. All measurements were made on the high resolution images.

## 3. Results

### 3.1. FPM background correction reduces non-uniformity in FPM reconstructions

The background correction approach was first evaluated using representative amplitude (Figure 2) and phase contrast (Figure 3) FPM reconstructions. In the raw FPM amplitude image, background non-uniformities were visible across the field of view, as shown in Figure 2(A). These variations were apparent following application of the 16 Colors LUT, shown in Figure 2(B), which revealed considerable background non-uniformity. A foreground mask successfully identified specimen-containing regions, allowing these regions to be excluded from the subsequent correction. These data are shown in Figure 2(C).

**Figure 2.**
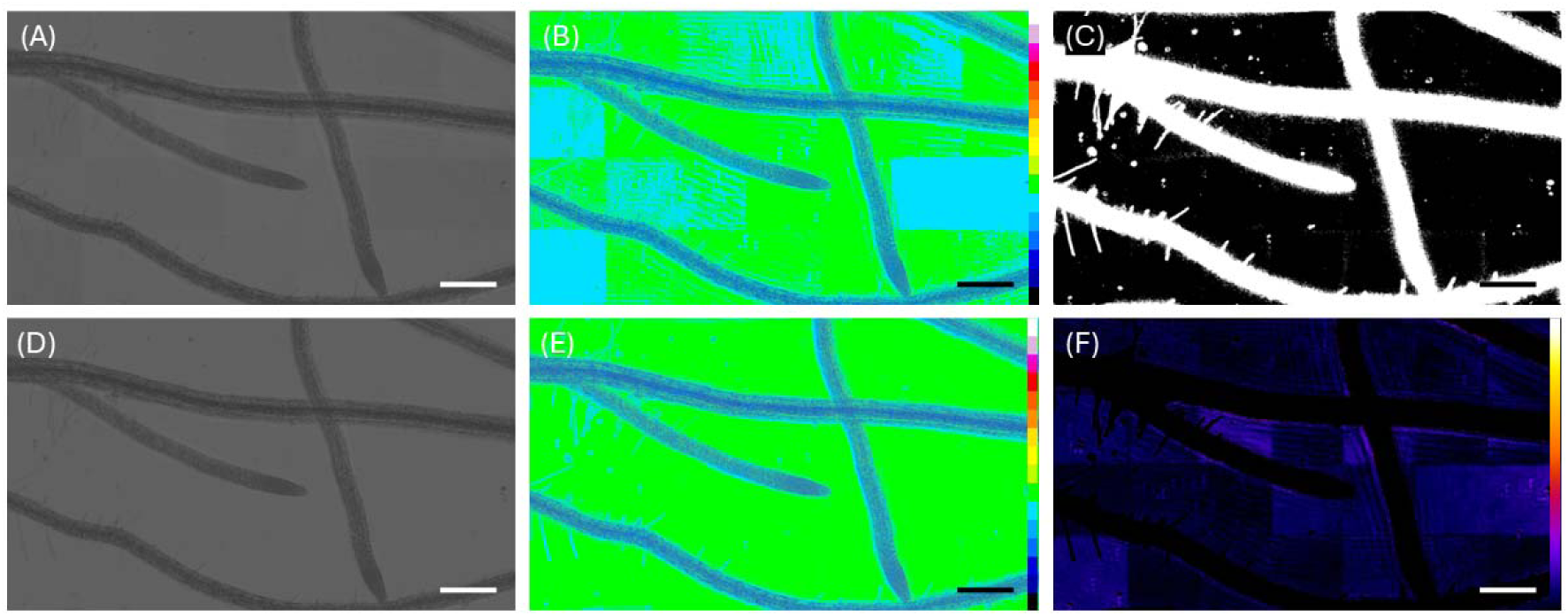
(A) Raw FPM amplitude image. (B) The same data shown in (A) displayed using the 16 Colors lookup table (LUT) to more clearly visualise non-uniformity in background arising from reconstruction artefacts. (C) Foreground mask identifying specimen regions excluded from the background correction. (D) Image data following background correction, showing a more uniform background while retaining the specimen information. (E) Corrected reconstruction shown in (D) displayed using the 16 Colors LUT. The same intensity limits were used in (B) and (E) to allow direct comparison before and after correction. (F) Absolute pixel-wise difference between the raw and corrected reconstructions, displayed using the Fire LUT from zero to the maximum difference. Changes in the image are restricted to the background, with no observed change within the masked specimen regions. Scale bars = 250 µm.

**Figure 3.**
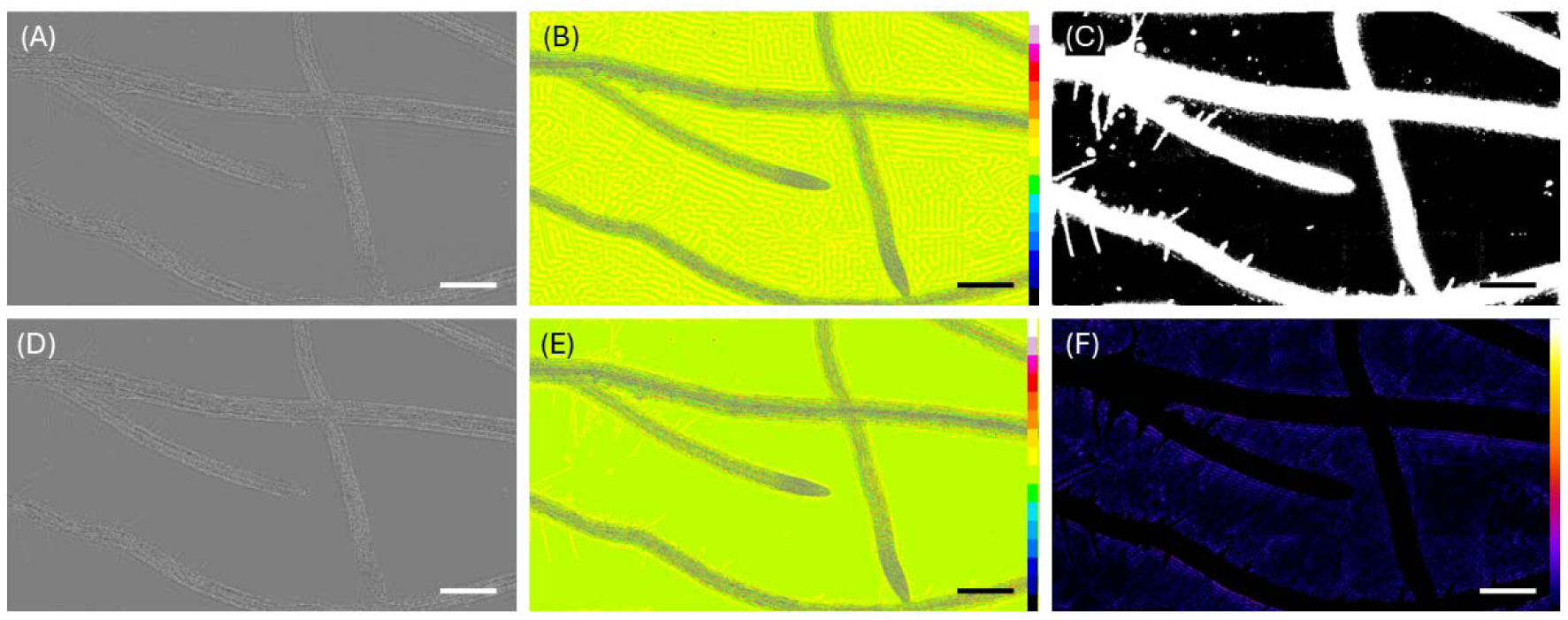
(A) Raw FPM phase image. (B) FPM phase image shown in (A) displayed using the 16 Colors lookup table (LUT) to more clearly show background non-uniformity. (C) Foreground mask identifying specimen regions excluded from background correction. (D) Image data following background correction, showing a more uniform background while retaining the specimen information. (E) Corrected reconstruction shown in (D) displayed using the 16 Colors LUT. (F) Absolute pixel-wise difference between the raw and corrected reconstructions, displayed using the Fire LUT from zero to the maximum difference. The corrected phase reconstruction shows a considerable reduction in background non-uniformity, with no change to the specimen information. Scale bars = 250 µm.

Application of the method substantially corrected the image background while preserving the appearance and intensity distribution of the specimen, as shown in Figure 2(D). This effect was especially evident when the corrected reconstruction was displayed using the 16 Colors LUT, presented in Figure 2(E), where background non-uniformity was considerably reduced. The absolute pixel-wise difference between the raw and corrected reconstructions, as shown in Figure 2(F), confirmed that changes were largely confined to the background with no visible change to the specimen.

A similar effect was observed in the reconstructed phase image. The uncorrected FPM phase estimate image contained background structure that was difficult to appreciate fully in the greyscale representation shown in Figure 3(A) but became visible following pseudo-colour display in Figure 3(B). Following foreground masking as shown in Figure 3(C), correction produced a much more uniform background while retaining the phase features associated with the specimen. These data are shown in Figures 3(D) and 3(E). Root hairs branching from the main root cannot be clearly seen in 3(B) because of the non-uniform background, but following background correction these are visible in 3(E). As with the amplitude reconstruction, the absolute difference image in Figure 3(F) confirmed that the correction modified only pixels outside the protected specimen mask, leaving protected specimen pixel values unchanged.

Mean execution times for the *A. thaliana* amplitude and phase reconstructions, each with a file size of 292 MB, were 42.16 ± 0.51 s and 40.35 ± 0.15 s, respectively (n=3 runs per image), including file input/output and diagnostic output generation.

Together, these results demonstrate that the method can selectively suppress background non-uniformity in both amplitude and phase FPM reconstructions without substantially altering specimen-associated information. The preservation of specimen features despite considerable modification of the background is consistent with the mask-based design of the correction, in which foreground pixels are explicitly protected from background correction.

### 3.2 FPM background correction is effective across different biological specimens

Background correction was effective across images containing different specimen types and patterns of background non-uniformities, as shown in Figure 4. In the raw images, shown in Figures 4(A), 4(C), 4(E), and 4(G) with the 16 Colors LUT applied, non-uniformities in the background presented as tile-like regions and periodic and aperiodic local changes in intensity. Following correction, these non-uniformities were reduced, producing a more uniform background in each dataset, as shown in the counterpart images Figures 4(B), 4(D), 4(F), and 4(H). Importantly, specimen features remained clearly visible after correction. These examples demonstrate that the correction is applicable to a range of biological samples and background patterns without obvious loss of specimen-associated information.

**Figure 4.**
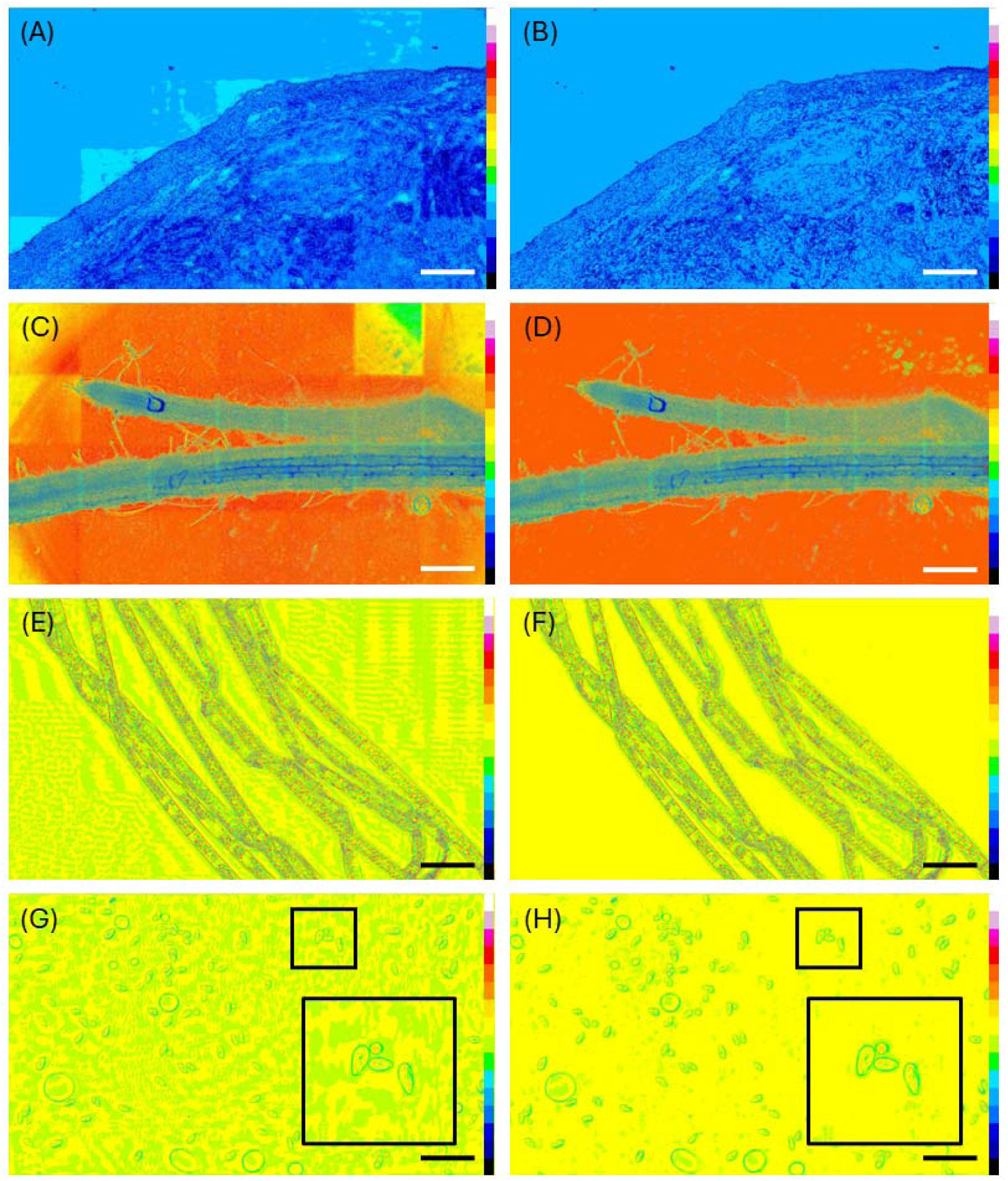
FPM background correction across multiple specimen types. Representative images are shown before (A, C, E, G) and after (B, D, F, H) background correction. (A, B) Human tonsil tissue stained with haematoxylin and eosin imaged with amplitude contrast; (C, D) *Nicotiana benthamiana* imaged with amplitude contrast; (E, F) *Spirogyra varians* imaged with phase contrast; and (G, H) *Tetrahymena pyriformis* with a silver nitrate stain imaged with phase contrast. Images were acquired in greyscale and are displayed using the 16 Colors lookup table (LUT) applied in FIJI to more clearly show background non-uniformities. Digital zoomed regions of interest (ROIs) are shown as insets in (G) and (H), highlighting the difference in background before and after correction. For each specimen, the same display range was used for the corresponding raw reconstructions and corrected images. Scale bars = 250 µm.

### 3.3 FPM background correction improves object detection without reducing image resolution

The effect of background correction on automated object detection was assessed using FPM amplitude and phase reconstructions of 5 µm silica beads, as shown in Figures 5 and 6, respectively. In both reconstructions, background non-uniformity hampered segmentation. Within the selected ROIs, FIJI Analyze Particles detected 251 objects in the raw amplitude FPM reconstruction and 13 objects in the raw phase contrast FPM reconstruction, despite each ROI containing only three silica beads. Following FPM background correction, the same analysis correctly identified the three beads in both reconstructions.

**Figure 5.**
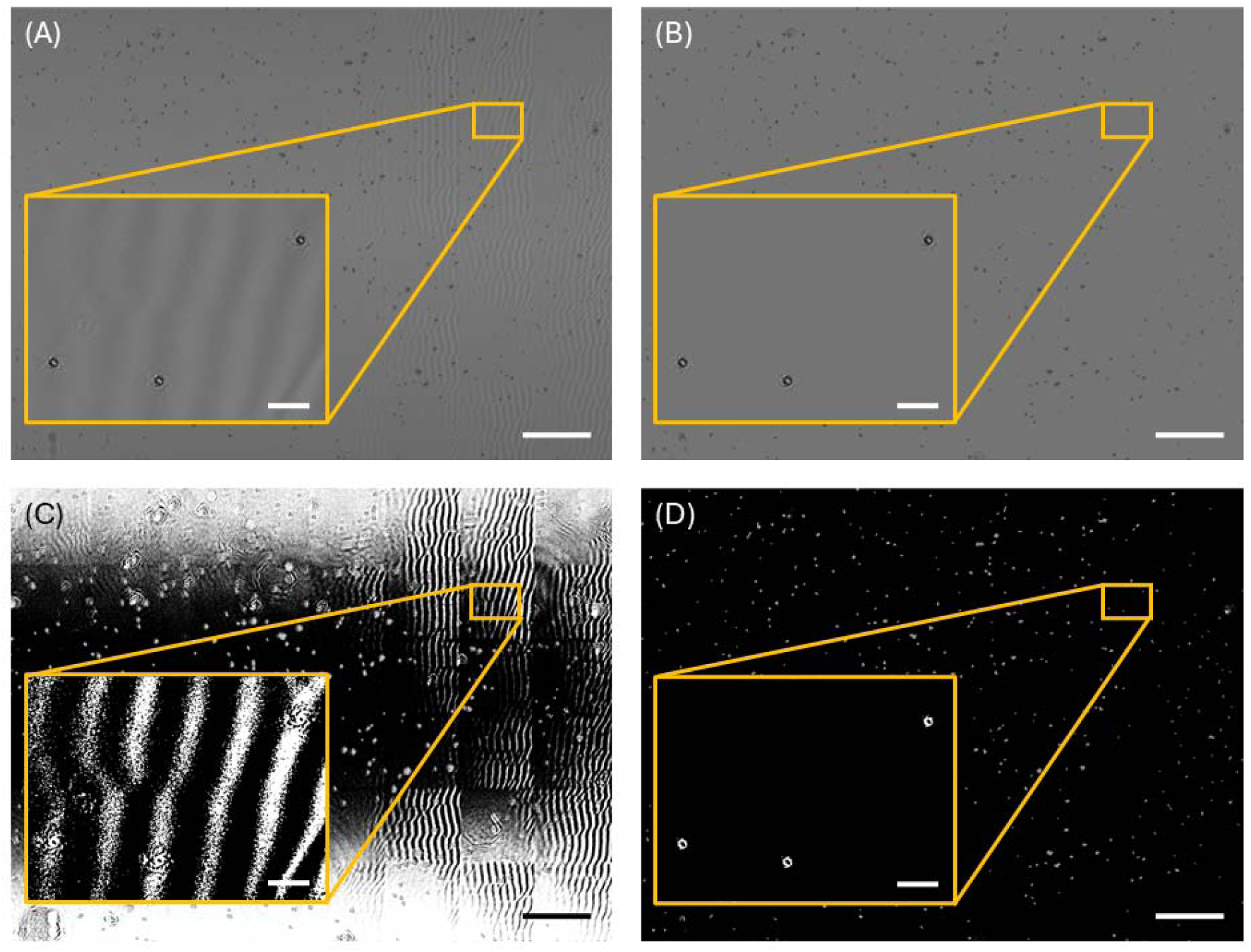
(A) Raw amplitude FPM reconstructed image of 5 µm silica beads mounted in Histomount. (B) Background corrected output of (A). In both (A) and (B) a digital region of interest (ROI) is highlighted with an orange box, and a digitally zoomed region is shown. In the ROI in (A), three beads are shown with a varying background; in (B), following FPM background correction, the same three beads are visible but with a homogeneous background. (C) Otsu thresholded version of (A). (D) Otsu thresholded version of (B). The same ROIs are highlighted, and digital zooms are shown. The background in the ROI of (C) obscures the thresholded beads and they cannot be seen. In the digital zoom of the ROI highlighted in (D), the thresholded particles are now clearly visible relative to the background. Using the Analyze Particles tool in FIJI, with settings of size (µm^2^): 0.8-Infinity and circularity=0-1, 251 objects are measured in the ROI in (C) compared to the three objects detected in the ROI in (D). Scale bars of the full images shown in (A-D) = 250 µm, scale bars of the digitally zoomed ROIs = 20 µm.

**Figure 6.**
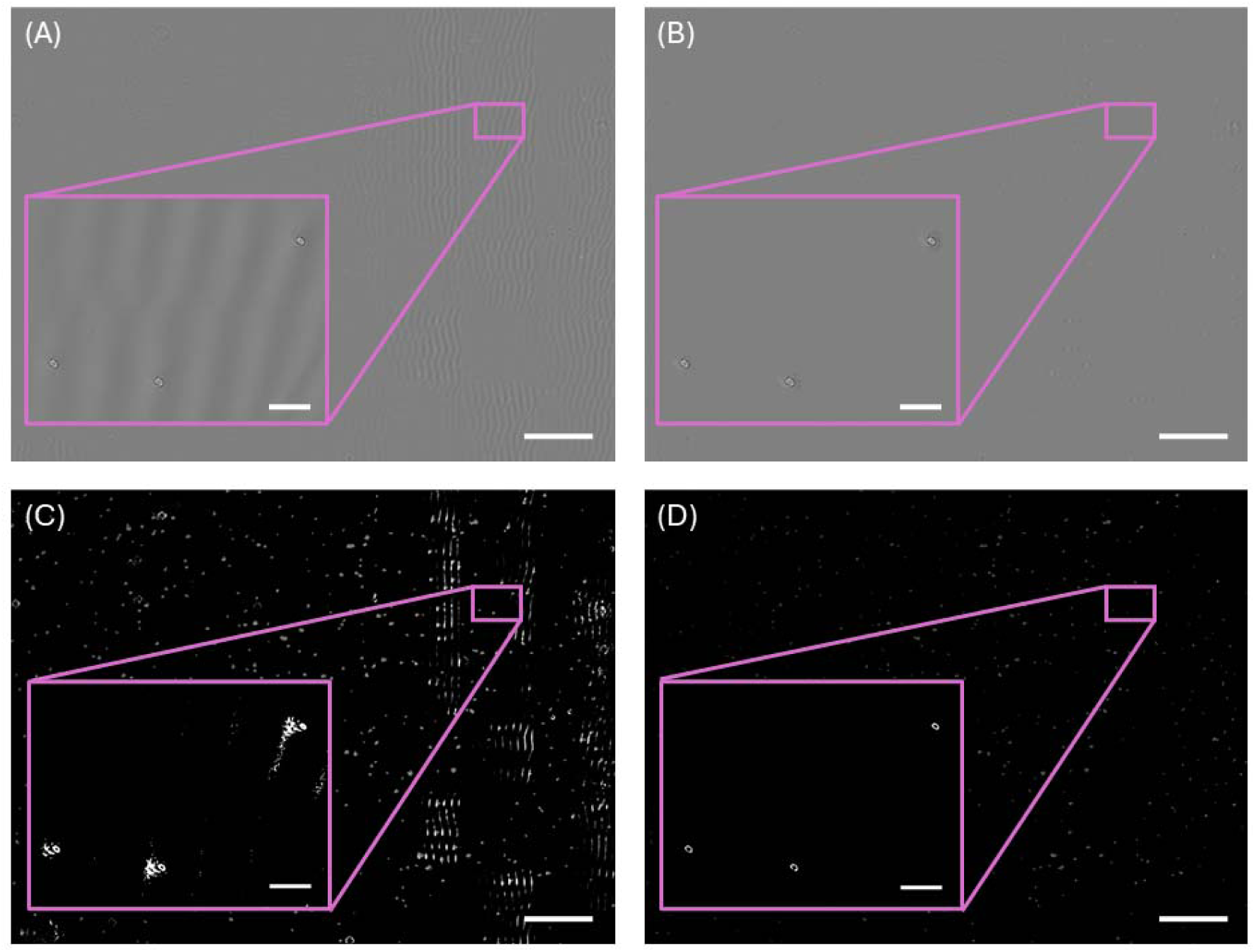
(A) Raw phase contrast FPM reconstructed image of 5 µm silica beads mounted in Histomount. (B) Background corrected output of (A). In both (A) and (B) a digital region of interest (ROI) is highlighted with a pink box, and a digitally zoomed region is shown. In the ROI in (A), three beads are shown with a varying background similar to that shown in Figure 5; in (B), following FPM background correction, the same three beads are visible but with a homogeneous background. (C) Otsu thresholded version of (A). (D) Otsu thresholded version of (B). The same ROIs are highlighted, and digital zooms are shown. The background in the ROI of (C) shows the thresholded beads but they are accompanied by background artefacts that surround each bead. In the digital zoom of the ROI highlighted in (D), the thresholded particles are now clearly visible relative to the background with no such artefacts. Using the Analyze Particles tool in FIJI, with settings of size (µm^2^): 0.8-Infinity and circularity=0-1, 13 objects are measured in the ROI in (C) compared to the three objects detected in the ROI in (D). Scale bars of the full images shown in (A-D) = 250 µm, scale bars of the digitally zoomed ROIs = 20 µm.

To determine whether background correction affected image resolution, image decorrelation analysis was performed on three corresponding ROIs in the raw and corrected reconstructions for each of four specimen types (Table 1). Mean resolution was 504 ± 34 nm before and 491 ± 32 nm after correction for *N. benthamiana*, 646 ± 63 nm and 592 ± 31 nm for tonsil, 644 ± 68 nm and 644 ± 68 nm for *S. varians*, and 710 ± 123 nm and 721 ± 109 nm for *Tetrahymena*, respectively. No statistically significant differences between paired raw and corrected measurements were detected for any of the four specimen types (two-tailed paired *t*-tests, *P* > 0.05 in all cases). Moreover, there was no consistent direction of change following correction: the mean decorrelation-based resolution decreased for *N. benthamiana* and tonsil tissue, was unchanged for *S. varians*, and increased slightly for *T*. *pyriformis*. These results indicate that background correction did not produce a significant or systematic loss of image resolution in the reconstructed images.

**Table 1.** Effect of background correction on image resolution. Image resolution was measured using image decorrelation analysis in n=3 corresponding ROIs before and after background correction for each specimen type. Resolution values are presented as mean ± standard deviation. No statistically significant differences were detected (*P* > 0.05 for all comparisons).

| Specimen | Raw image resolution (nm) | Corrected image resolution (nm) | P value |
| --- | --- | --- | --- |
| *N. benthamiana* | $504 \pm 34$ | $491 \pm 32$ | 0.063 |
| Tonsil tissue | $646 \pm 63$ | $592 \pm 31$ | 0.34 |
| *S. varians* | $644 \pm 68$ | $644 \pm 68$ | 1 |
| *T. pyriformis* | $710 \pm 123$ | $721 \pm 109$ | 0.75 |

The Fourier spectra of the original and background corrected *N. benthamiana* and *T. pyriformis* reconstructions are shown in Figure S1. The FFTs showed visually similar overall distributions and spatial frequency extents. These qualitative observations were consistent with the absence of a systematic loss of resolution in the decorrelation analysis.

### 3.4 Comparison with pseudo flat field correction

Pseudo flat field correction was used to compare background removal and preservation of specimen structure. A raw FPM amplitude reconstructed image of *Arabidopsis thaliana* is shown in Figure S2(A), with the background corrected image shown in Figure S2(B). Figures S2(C)-S2(F) show the result of S2(A) corrected using pseudo flat field correction with blurring radii of 50, 150, 250 and 350 pixels, respectively.

At a blurring radius of 50 pixels, pseudo flat field correction reduced residual background non-uniformity to 0.59%, but substantially altered the specimen, producing a structural error of 6.13%. In contrast, FPM background correction achieved a comparable background uniformity of 0.61% while preserving specimen structure, with a structural error of only 0.04%. Increasing the blurring radius reduced the change to the specimen information, but at the expense of poorer background correction, as shown in Figure S2(G). Overall, FPM background correction achieved comparable background uniformity while producing substantially less alteration of specimen information than pseudo flat field correction.

## 4 Discussion

This study demonstrates that background non-uniformity in FPM reconstructions can be reduced while preserving specimen-level information. The method was effective across amplitude and phase reconstructions and multiple biological specimen types, and improved object detection without a loss of image resolution. Importantly, this FPM background correction approach achieved similar background uniformity to pseudo flat field correction while producing substantially less structural change to the specimen. Together, these findings show that correction of the image background can improve the suitability of FPM reconstructions for quantitative analysis while preserving the specimen information required for accurate measurement.

Background non-uniformity is important when FPM images are used for quantitative analysis rather than visual interpretation alone. Variation in the background can alter the separation between specimen and background intensities across a large field of view, potentially affecting threshold-based segmentation and subsequent measurements.^16,17^ This was demonstrated directly using the silica bead data, where removal of background variation allowed the same thresholding and particle analysis to identify the expected objects correctly. The improvement therefore resulted not from enhancement of the objects themselves, but from removal of background variation that interfered with their segmentation. This demonstrates that background artefacts that may be relatively inconsequential for visual interpretation of FPM images can nevertheless have substantial consequences for downstream quantitative analysis.

The implications of improved background uniformity extend beyond individual image-analysis tasks. A major advantage of FPM is its ability to combine high resolution with a large imaging area, making it attractive for applications requiring quantitative analysis of large numbers of cells, tissue regions or other structures within a single dataset.^1,2^ As FPM technologies develop towards larger fields of view, higher throughput and increasingly automated acquisition and analysis^4,6,29^, consistency across reconstructed images becomes increasingly important. Background variation may otherwise introduce unwanted differences in segmentation or measurement, limiting comparisons across an extended specimen. Specimen-preserving background correction could therefore support applications such as large-area tissue imaging, digital pathology, cell phenotyping and high-content imaging, where quantitative measurements must remain comparable across large fields of view. More broadly, these findings suggest that background uniformity should be considered alongside image resolution, field of view and imaging throughput when evaluating FPM systems intended for quantitative applications. This consideration may become increasingly important as FPM is increasingly integrated with automated and machine-learning-based analysis, where systematic background variation could introduce unwanted variability into downstream classification or measurement.^30^

For these applications, however, improving background uniformity is only useful if the correction itself does not compromise the specimen information to be analysed. The decorrelation-based resolution measurements (Table 1), together with the visually similar spatial-frequency extent before and after correction (Figure S1), support the preservation of image detail. The FFTs provide qualitative supporting evidence, although they include contributions from both specimen and background and do not establish that individual frequency components remain unchanged. While appropriate for many applications, modification of specimen pixels is potentially problematic when the resulting image will subsequently be quantified, because the correction itself may alter the information being measured. The comparison with pseudo flat field correction^28^ demonstrates this distinction, where achieving a similar degree of background uniformity resulted in a change to the specimen-level information. The advantage of the present approach is therefore not simply improved background uniformity, but the separation of background correction from modification of the specimen itself. It is important to distinguish preservation of specimen pixel values from removal of all background-related artefacts affecting the specimen. Spatial variations arising during acquisition or reconstruction may extend into specimen-containing regions, even where they are less readily visible than in the surrounding background. The present method does not attempt to remove these contributions from protected pixels. Instead, it avoids introducing further changes to specimen intensities while improving the uniformity of the surrounding background. Correction of artefacts within specimen regions would require a different approach, potentially involving modelling of the background signal during acquisition or reconstruction.

This distinction is relevant to FPM, where background artefacts may originate from the acquired image data as well as from illumination, optical and reconstruction effects^9–12,31^. Previous approaches have therefore sought to address background interference at an earlier stage of the FPM pipeline. Hou *et al.* incorporated adaptive separation of background interference into the reconstruction model^10^, while Claveau *et al.* suppressed background signal and associated noise in the acquired dark-field images before their incorporation into iterative reconstruction.^9^ The present approach is complementary: rather than attempting to identify or model the source of imaging or reconstruction artefacts, it can be applied directly to the reconstructed image data.

There are limitations to this approach. Its performance depends on accurate identification of the specimen, and features that are not incorporated into the protected mask could be treated as background, while background incorrectly classified as specimen will remain uncorrected. The method is therefore most appropriate for images in which specimen-containing and background regions can be reliably distinguished. Images in which the specimen occupies most of the field of view, or in which weak specimen features cannot readily be separated from the background, may prove more challenging. In addition, the present implementation replaces background pixels with a uniform median value. This is advantageous where a homogeneous background is required for subsequent segmentation but may not be appropriate where intensity information within nominal background regions is itself of interest. For weakly absorbing specimens, phase reconstructions may provide greater contrast than amplitude reconstructions and could therefore offer a more reliable basis for specimen mask generation. Since both amplitude and phase reconstructions are available, the use of phase-derived or combined amplitude-phase masks may improve the identification of weak specimen features.

The quantitative evaluation was also limited to selected analysis tasks and datasets. The examples presented here demonstrate that background variation can substantially affect automated analysis and that effective whole-image correction can alter specimen information, but they do not constitute a comprehensive comparison with all available background-correction or segmentation approaches. Future work could evaluate the method across larger datasets, different FPM systems and reconstruction methods, and additional quantitative tasks including measurements of object area, morphology and intensity. Alternative implementations could also estimate a smoothly varying corrected background while retaining the central principle of protecting specimen-associated pixels from modification. Evaluation in emerging high-throughput and automated FPM platforms would be valuable in establishing the extent to which improved background uniformity increases measurement consistency across large datasets.

A potential extension of this approach is its application to other forms of spatial background variation in light microscopy. For example, large-area images assembled by tiling or stitching can contain tile-to-tile intensity differences, illumination gradients, and visible boundaries between adjacent fields which may similarly interfere with segmentation and quantitative analysis.^32^ Because the present method identifies specimen-containing regions and restricts correction to the surrounding background, it could potentially be adapted to reduce such artefacts without applying intensity transformations directly to the specimen. However, this remains to be established experimentally, and the current implementation was developed and evaluated specifically for background non-uniformity in FPM reconstructions. Evaluation across other microscopy modalities will therefore be required to determine whether this approach can provide a more general method for background correction.

These results demonstrate that background non-uniformity in reconstructed FPM images should be considered not only in terms of image appearance, but also in terms of its consequences for quantitative analysis. As FPM develops towards larger fields of view, higher throughput and greater integration with automated analysis, maintaining consistent image properties across the reconstructed field will become increasingly important. The specimen-preserving approach presented here reduces background non-uniformity without substantially altering specimen information or image resolution, providing a simple route towards more robust quantitative analysis of FPM image data.

## Data availability statement

The research data underpinning this publication can be accessed at https://doi.org/10.15129/02e4cc83-bedb-4005-9ca3-ca511a9013fc (DOI to be minted upon publication). The Python code for FPM background correction together with instructions for use and installation are available at https://github.com/gailmcconnell/FPM-Background-Correction.

---

**Figure S1.**
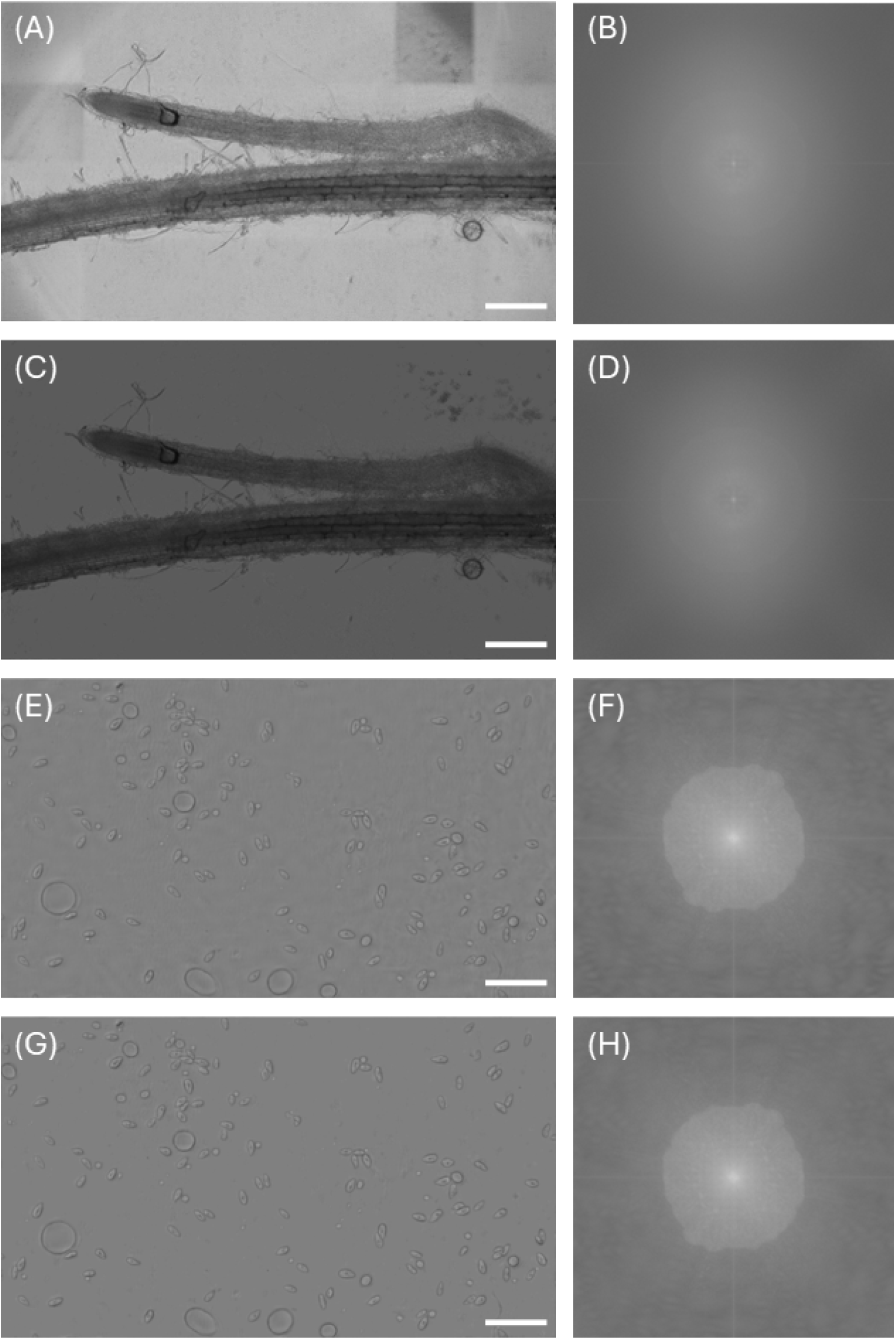
Spatial frequency content of FPM reconstructions before and after background correction. (A, C) Original and background corrected amplitude reconstructions of *Nicotiana benthamiana*, corresponding to Figures 4(C) and 4(D). (B) and (D) show two-dimensional fast Fourier transforms (FFTs) of the images in (A) and (C), respectively. (E) and (G) are duplicates of Figure 4(G) and 4(H), showing original and background corrected phase reconstructions of *Tetrahymena pyriformis*, and the corresponding FFTs are shown in (F) and (H) here. For both specimens, the overall distribution and extent of the spatial frequency content remain visually similar before and after background correction. Scale bars = 250 µm.

**Figure S2.**
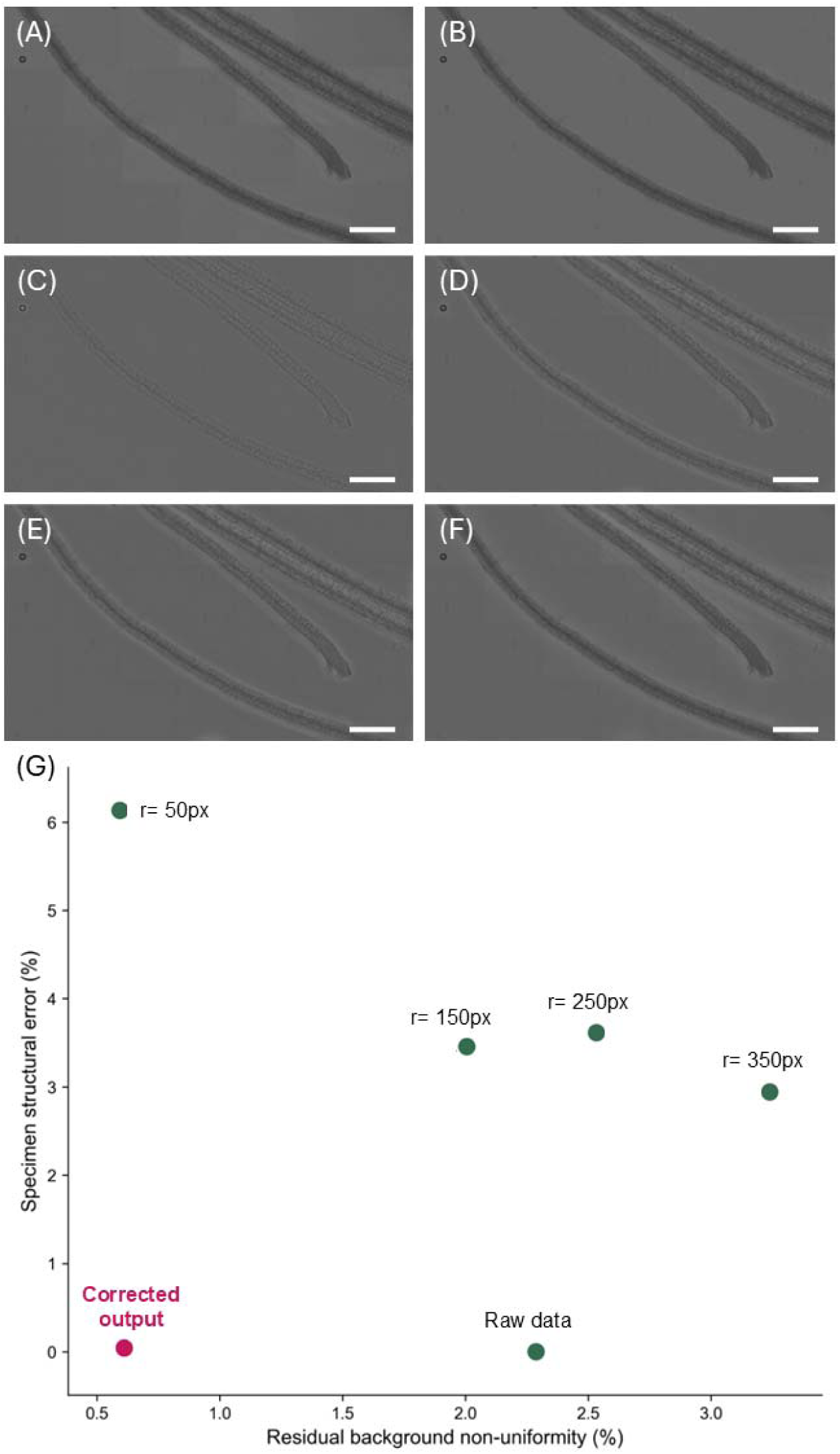
(A) Raw amplitude FPM reconstructed image of *Arabidopsis thaliana*. (B) Background corrected reconstruction of (A). (C-F) Pseudo flat field correction applied to (A) with blurring radii of (C) 50, (D) 150, (E) 250 and (F) 350 pixels. In (C-F), all changes to the specimen are visible, with correction of the background varying depending on the blurring radius. Scale bars = 250 µm. (G) Residual background non-uniformity plotted against specimen structural error for the raw data, FPM background corrected output, and pseudo flat field corrections for the different blurring radii. Residual background non-uniformity was calculated as the coefficient of variation of background tile intensities, while specimen structural error represents the normalised root mean square error relative to the raw image after allowing for differences in overall intensity and offset. A blurring radius of r=50 pixels with pseudo flat field correction gave low background non-uniformity (0.59%) but a structural error of 6.13%, while FPM background correction gave a similar background non-uniformity (0.61%) with a structural error of only 0.04%.

